# Nucleoid Compaction Influences Carboxysome Localization and Dynamics in *Synechococcus elongatus* PCC 7942

**DOI:** 10.1101/2025.04.21.649805

**Authors:** Claire E. Dudley, Christopher A. Azaldegui, Daniel J. Foust, Olivia LaCommare, Julie S. Biteen, Anthony G. Vecchiarelli

## Abstract

The bacterial nucleoid is not just a genetic repository—it serves as a dynamic scaffold for spatially organizing key cellular components. ParA-family ATPases exploit this nucleoid matrix to position a wide range of cargos, yet how nucleoid compaction influences these positioning reactions remains poorly understood. We previously characterized the Maintenance of Carboxysome Distribution (Mcd) system in the cyanobacterium *Synechococcus elongatus* PCC 7942, where the ParA-like ATPase McdA binds the nucleoid and interacts with its partner protein, McdB, to generate dynamic gradients that distribute carboxysomes for optimal carbon fixation. Here, we investigate how nucleoid compaction impacts carboxysome positioning, particularly during metabolic dormancy when McdAB activity is downregulated. We demonstrate that a compacted nucleoid maintains carboxysome organization in the absence of active McdAB-driven positioning. This finding reveals that the nucleoid is not merely a passive substrate for positioning, but an active player in spatial organization. Given the widespread role of ParA-family ATPases in the positioning of diverse cellular cargos, our study suggests that nucleoid compaction state is a fundamental, yet underappreciated, determinant of mesoscale organization across bacteria.

## INTRODUCTION

Bacteria have evolved diverse strategies to spatially organize intracellular components. The ParA family of positioning ATPases is widespread across the bacterial kingdom, regulating a variety of cellular cargos, including chromosomes, plasmids, protein-based complexes, and organelles ^1–6^. Regardless of cargo type, the bacterial nucleoid serves as the positioning matrix. Upon ATP binding, ParA ATPases dimerize, enabling them to bind nucleoid DNA ^7–11^. This dimerization also creates a binding site for an ATPase-activating partner protein that associates with the cargo, locally stimulating ParA ATPase activity and promoting its release from the nucleoid ^6,12,13^. This dynamic interplay generates ParA gradients that distribute cargos along the nucleoid via a diffusion-ratchet mechanism ^14,15^. Despite current models treating the nucleoid as a passive matrix for cargo positioning ^2^, the nucleoid is highly dynamic, continuously compacting and decompacting in response to environmental cues ^16–20^. It remains unclear how these fluctuations in compaction state influence ParA-mediated cargo organization in bacterial cells.

Bacterial Microcompartments (BMCs) are protein-based organelles that encapsulate key metabolic processes ^21,22^. A recent bioinformatics survey identified 68 BMC types across 45 bacterial phyla ^22^. Despite their prevalence and metabolic significance, little is known about how BMCs are spatially regulated within the cell. The carboxysome, a carbon-fixing BMC, is the best-studied example of BMC subcellular organization ^14,23–28^. Carboxysomes are widespread among cyanobacteria and some chemoautotrophs, contributing signifcantly to global CO_2_ fixation ^14,29^, making them of significant ecological and biotechnological interest.

In the model rod-shaped β-cyanobacterium *Synechococcus elongatus* PCC 7942 (*S. elongatus* hereafter), a two-protein system ensures the even distribution of carboxysomes along the cell length during exponential growth under optimal conditions ^3,14^. The maintenance of carboxysome distribution protein A (McdA) is a ParA family ATPase that binds nucleoid DNA in its ATP-bound dimeric form ^3,14,25^. Its partner protein, McdB, localizes to carboxysomes and drives McdA oscillations which facilitate the equidistant positioning of carboxysomes ^14,24^. In the absence of McdAB, carboxysomes mislocalize into nucleoid-excluded clusters, leading to their rapid loss from the cell population and slower autotrophic growth ^14,24^.

Here, we investigate the role of the nucleoid and its compaction state in the spatial organization of carboxysomes. Under cellular stress, when the McdAB system is downregulated, we find that the compacted nucleoid helps maintain carboxysome organization. Additionally, our findings indicate that nucleoid compaction influences carboxysome dynamics within the cell. We propose that nucleoid compaction plays a crucial role in organizing mesoscale cellular cargos in bacteria. Specifically, we show that a compacted nucleoid can anchor distributed carboxysomes in place, even in the absence of the McdAB system. These findings have broad implications in establishing the nucleoid as an active participant—rather than a passive scaffold—in intracellular organization. They suggest that global changes in nucleoid architecture can directly influence the spatial dynamics of all cellular cargos organized by ParA family ATPases, fundamentally reshaping our understanding of bacterial cell organization and its adaptability under stress.

## RESULTS

### Carboxysomes remain distributed in stationary phase cells

We have shown that carboxysomes in *S. elongatus* are distributed along the cell length by the McdAB system in exponential phase while under optimal growth conditions ^14,24^. Cameron et al. showed that carboxysomes in stationary phase cells (∼ 30 days on solid BG11 media) remain distributed ^30^. Intriguingly, the presence of chlorophyll indicated these carboxysome-containing cells at stationary phase remain photosynthetically active and viable; as dead cells were devoid of both chlorophyll and carboxysomes. The results suggest that carboxysome maintenance is critical to cellular fitness during extended periods of nutrient limitation. However, it remains to be determined if and how the McdAB system maintains carboxysome organization during nutrient stress.

We first set out to determine how carboxysome organization is influenced when cells transition from exponential growth to stationary phase in batch culture. We used a growth curve to determine the exponential and stationary phases of *S. elongatus* growth under our standard conditions (Figure S1A). To image carboxysomes, the fluorescent protein monomeric Turquoise2 (mTQ) was fused to the C-terminus of the small subunit of Rubisco (RbcS) to make RbcS-mTQ (see Materials and Methods for strain construction details). *RbcS-mTQ* was expressed using a second copy of its native promoter (inserted at neutral site 1) in addition to wild-type *rbcS* at its native locus in wild-type cells. Growth of the fluorescently labeled strains was similar to that of wildtype (WT) (Figure S1A). As shown previously, RbcS-mTQ-labeled carboxysomes are distributed down the long axis of *S. elongatus* cells during exponential growth (Figure 1A). In stationary phase, viable cells with a chlorophyll signal also had carboxysomes distributed along the cell length. However, carboxysomes were more tightly packed along the medial axis of the cell (Figure 1BC), with a loss in the hexagonal packing of carboxysomes found in exponential cells (Figure 1A, arrow) ^14^. The tight linear arrangement suggested that carboxysome diffusion was also restricted in stationary phase cells.

**Figure 1.**
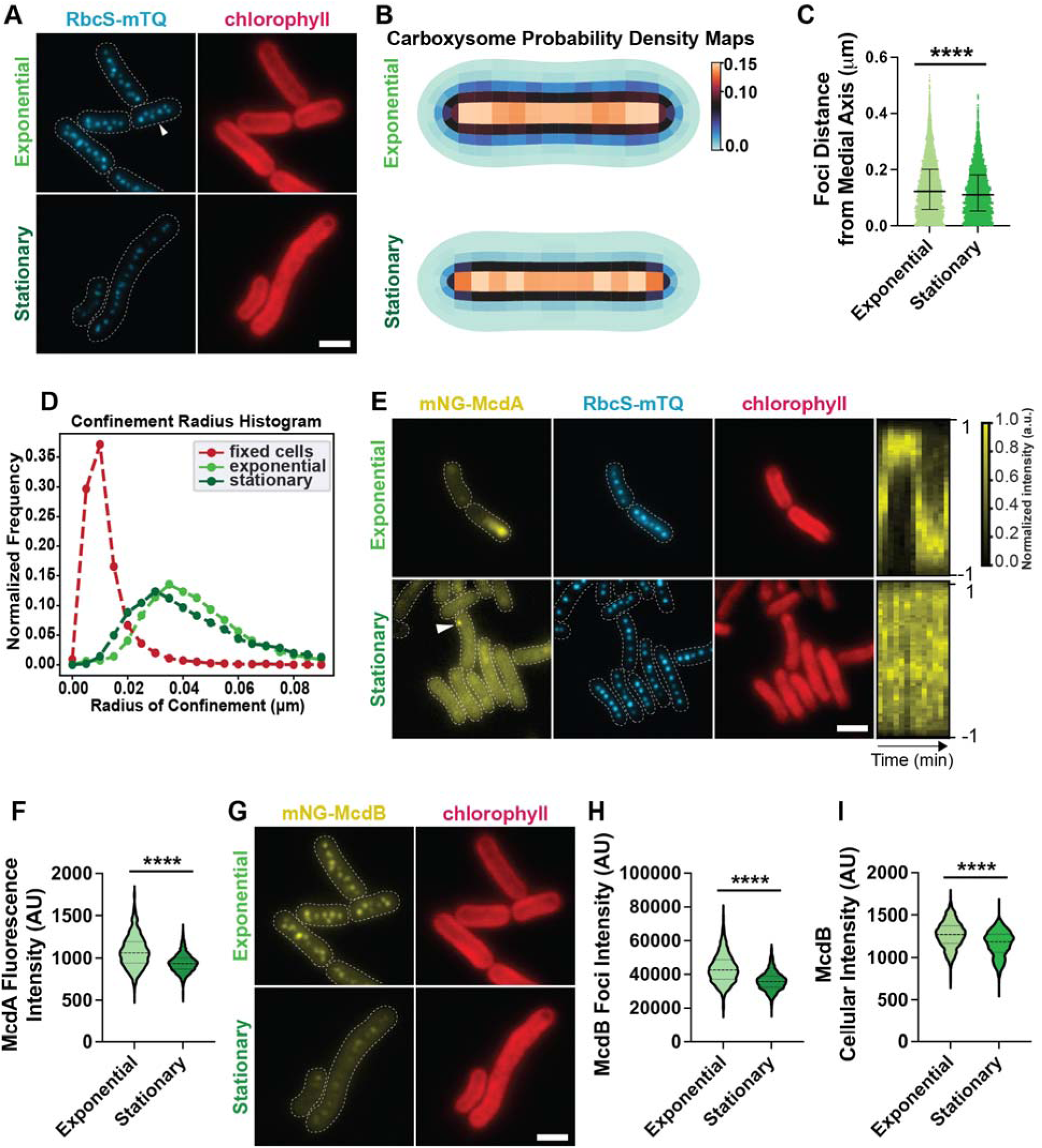
Carboxysomes are positioned in living stationary phase cells. **(A)** Representative microscopy images of carboxysomes in exponential and stationary phase cells. Carboxysomes are in cyan and the cell boundary from the phase contrast channel is a dashed white line. White arrow highlights hexagonal packing of carboxysomes in exponential phase cells. The chlorophyll channel is in red. **(B)** Carboxysome probability density maps from cells in each growth phase. Exponential, n = 1662 cells; stationary, n = 1401 cells. **(C)** Carboxysome foci distance from the medial axis. Distances of carboxysome foci from the long axis of the cell, computed using particle mapping code (see Methods). Significance from Mann-Whitney test, P < 0.0001, (exponential) n = 3 biological replicates and (stationary) n = 2 biological replicates, >200 cells each. **(D)** Confinement radius histogram illustrating carboxysome foci confinement radii in exponential, stationary, and fixed cells based on TrackMate analysis. **(E)** Microscopy images of mNG-McdA and carboxysome foci in exponential and stationary phase cells. Dotted white line shows cells’ boundaries from phase contrast. Chlorophyll channel is in red. Kymographs of mNG-McdA dynamics from a representative cell in exponential and stationary phase are also provided. Short-axis is time, 0 to 60 minutes, the long-axis is cell length. **(F)** mNG-McdA whole cell fluorescence intensity normalized by cell length. Significance from Mann-Whitney test, P < 0.0001, (exponential) n = 3 biological replicates and (stationary) n = 2 biological replicates, >200 cells each. **(G)** Microscopy images of mNG-McdB in exponential and stationary phase cells. Chlorophyll channel is in red. **(H)** mNG-McdB foci intensity. Significance from Mann-Whitney test, P < 0.0001, (exponential) n = 3 biological replicates and (stationary) n = 2 biological replicates, >200 cells each. **(I)** Whole cell McdB fluorescence intensity normalized by cell length. Significance from Mann-Whitney test, P < 0.0001, (exponential) n = 3 biological replicates and (stationary) n = 2 biological replicates, >200 cells each. (scale bar 2 μm).

Consistently, an analysis on the confinement radius of carboxysome diffusion found that carboxysome localization and movement are both restricted in stationary phase cells (Figure 1D, **Video 1, Video 2**).

Carboxysome foci in stationary phase cells displayed a subtle decrease (∼8%) in RbcS-mTQ intensity (Figure S1B) and slightly tighter spacing (Figure S1C) relative to exponential phase. However, carboxysome copy number per cell was similar to that of exponential phase cells (Figure S1D). Together, the data suggest that carboxysome composition likely changes as cell populations age, but the carboxysome distribution is maintained in stationary phase cells.

### Localization of the McdAB system is perturbed during stationary phase

We have previously shown that the McdAB system is responsible for distributing carboxysomes in *S. elongatus* ^14,24^. Without the McdAB system, mispositioned carboxysomes form nucleoid-excluded clusters ^14,25^. Given our data showing that carboxysomes remain distributed in stationary phase, we set out to determine if the McdAB system was responsible.

To simultaneously image McdA or McdB in our carboxysome reporter strain, McdA or McdB was N-terminally fused to the fluorescent protein monomeric NeonGreen (mNG) ^31^. We have previously shown that mNG-McdA and mNG-McdB are functional for carboxysome positioning when expressed as the only copy at their native locus ^14^. Growth of these fluorescently labeled strains was similar to that of WT (Figure S1A). We also performed phase-contrast imaging to monitor changes in cell morphology.

As shown previously, McdA oscillates from pole to pole on the nucleoid in *S. elongatus* cells during exponential growth (Figure 1E, Figure S1E, **Video 3**). Intriguingly, robust McdA oscillation was lost in stationary phase cells (Figure 1E, Figure S1F, **Video 4**). Instead, most stationary cells with a chlorophyll signal displayed diffuse McdA in the cytoplasm, despite carboxysomes remaining distributed across the cell length. A few cells also contained bright non-oscillatory McdA foci that did not colocalize with carboxysomes (Figure 1E, arrow). We have shown previously that the mNG-McdA signal intensity is proportional to McdA levels in the cell ^14,25^. Quantification of whole cell McdA fluorescence intensity, normalized by cell length, found that the median McdA level in stationary phase cells only decreased ∼ 14% compared to exponential phase cells (Figure 1F). The data suggest that the McdA remaining in stationary phase cells is no longer competent for dynamic nucleoid binding and oscillation, yet carboxysomes remain distributed.

The intensity of mNG-McdB associated with carboxysomes decreased significantly in stationary phase cells (Figure 1GH). Yet, quantification of whole cell mNG-McdB fluorescence intensity, normalized by cell length, found that the median McdB level only decreased ∼ 8% compared to exponential phase cells (Figure 1I). Together, our data show that while the cellular levels only decline slightly for both McdA and McdB, McdA no longer associates with the nucleoid, and McdB no longer colocalizes with carboxysomes in the stationary phase. The evidence suggests that the McdAB system is not the major driver of carboxysome distribution in stationary phase cells, which led us to consider other factors.

### Phosphate depletion compacts the nucleoid and downregulates McdAB, yet carboxysomes remain distributed

The nucleoid serves as a critical matrix for positioning diverse cellular cargos by ParA family ATPases, including chromosomes, plasmids, and an array of protein-based organelles, many of which are fundamental to cell survival and pathogenesis ^2,32,33^. Yet it remains to be determined how the compaction state of the nucleoid influences the subcellular organization of these mesoscale cargos. Consistent with other bacterial species ^17,18,34^, the nucleoid of *S. elongatus* significantly compacts in stationary phase (Figure 2AB). Nucleoid compaction correlated with carboxysomes aligning down the short axis of stationary-phase cells (Figure 2A). Together, the data suggest that the compacted nucleoid holds distributed carboxysomes in place in stationary phase cells when the McdAB system is downregulated.

**Figure 2.**
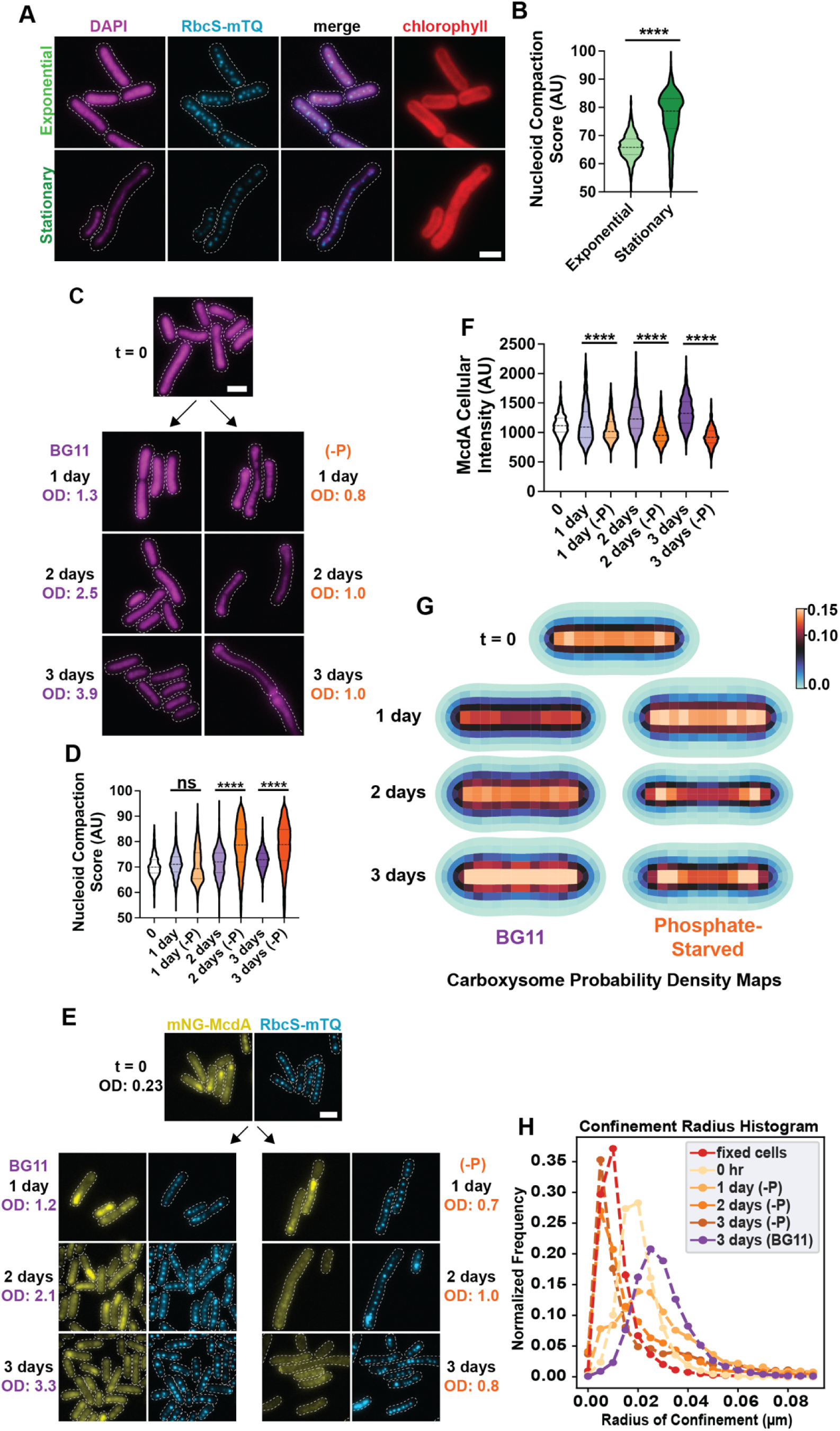
Phosphate deprivation induces nucleoid compaction and downregulation of McdAB. **(A)** Microscopy images of DAPI-stained nucleoids (magenta) in exponential and stationary phase cells. Dotted white line shows cell boundaries from phase contrast. Carboxysomes are in cyan and the merge shows carboxysome and DAPI channels. Red is the chlorophyll channel. **(B)** Nucleoid compaction score in arbitrary units (see Methods). Significance from Mann-Whitney test, P < 0.0001, (exponential) n = 6 biological replicates and (stationary) n = 5 biological replicates, >200 cells per replicate. **(C)** Microscopy images of DAPI-stained nucleoids in cells grown in BG11 or phosphate-depleted (-P) media. Dotted white line shows cell boundaries from phase contrast. **(D)** Nucleoid compaction score in arbitrary units in non-starved and phosphate-limited (-P) cells. Significance from Kruskall-Wallis and Dunn’s multiple comparisons, P < 0.0001. Biological replicates, (BG11) n = 4 and (-P) n = 6, > 200 cells per replicate. **(E)** Microscopy images of mNG-McdA and carboxysome foci in cells grown in BG11 and phosphate-limited BG11 (-P) media over a 3-day time course. Dotted white line shows cell boundaries from phase contrast. **(F)** mNG-McdA whole cell fluorescence intensity normalized by cell length. Significance from Kruskall-Wallis and Dunn’s multiple comparisons, P < 0.0001. Biological replicates, (BG11) n = 2 and (-P) n = 3, > 300 cells per replicate. **(G)** Carboxysome probability density map from cells grown in BG11 and phosphate deprivation conditions. t = 0, n = 1037 cells; 1 day: n = 629 cells (BG11), n = 1216 cells (-P); 2 days: n = 1079 cells (BG11), n = 1490 cells (-P); 3 days: n = 1857 cells (BG11), n = 1317 cells (-P). **(H)** Confinement radius histogram illustrating carboxysome diffusion radii in cells grown in the conditions indicated. (scale bar 2 μm).

Based on this finding, we set out to determine the specific conditions in stationary phase driving nucleoid compaction. It has been shown previously that phosphate depletion induces nucleoid compaction in *S. elongatus* ^16,35^. Indeed, cells grown in phosphate-free BG11 media exhibited significant nucleoid compaction, compared to cells grown in phosphate-sufficient media (Figure 2CD). The degree of nucleoid compaction after two days of phosphate starvation was comparable to that found in stationary phase cells (see Figure 2B). Phosphate starvation also halted cell growth (Figure S2A). The data suggest that phosphate depletion serves as a key driver of nucleoid compaction and the resulting alignment of carboxysomes in stationary phase cells.

Observing these significant changes in the topology of the positioning matrix under phosphate depletion led us to consider how McdA oscillation and carboxysome distribution are affected. We found that McdA oscillations dampened after one day of phosphate depletion and were largely undetectable after three days (Figure 2E, Figure S2B). McdA cellular levels were also lower in phosphate-depleted cells compared to cells grown in BG11 (Figure 2F). Despite the decrease in McdA levels and loss of McdA oscillations, carboxysome foci number, intensity, and spacing did not drastically change (Figure S2C-E). Also, McdB colocalization and intensity on carboxysomes only changed moderately over the 3-day phosphate starvation period (Figure S2FG). However, as in stationary phase cells, carboxysomes were better aligned along the medial axis in phosphate-starved cells (Figure 2G).

Interestingly, in phosphate-deprived cells showing areas of both nucleoid expansion and compaction, carboxysomes were linearly distributed along the compressed stretches of nucleoid and clustered in areas of expansion (Figure S2H). Consistent with this observation, the degree of nucleoid compaction in phosphate-deprived cells also correlated with restricted carboxysome diffusion in the cell (Figure 2H). After two days of phosphate deprivation, carboxysomes were essentially static in the cell. Together, we conclude that phosphate depletion results in the downregulation of the McdAB system and nucleoid compaction, which immobilizes carboxysomes and maintains their linear distribution in the absence of active positioning.

### McdA is sequestered, carboxysomes cluster, and the nucleoid moderately compacts in prolonged darkness

The circadian clock of *S. elongatus* has been shown to control nucleoid compaction— compacting at dusk and expanding through the night, effectively regulating gene expression through these physical changes in chromosome structure ^36^. However, the effects of prolonged light limitation due to high cell densities at stationary phase in batch culture or in the environment have not been investigated. Therefore, to determine if light limitation also influences nucleoid compaction, we incubated cells in continuous darkness for five days. During this time, growth halted, showing that the cells were metabolically dormant (Figure S3A). After one day in darkness, the nucleoid did not dramatically compact compared to cells grown in light (Figure 3AB). After two days, nucleoid compaction was only moderate, compared to the compaction we observed after 2 days of phosphate deprivation. Even after five days in darkness, the degree of compaction was only half of that found in stationary phase cells or phosphate-starved cells (see Figure 2B and 2D). Reintroduction into continuous light for one day expanded the nucleoid (Figure 3AB), with a recovery in cell growth as well (Figure S3A).

**Figure 3.**
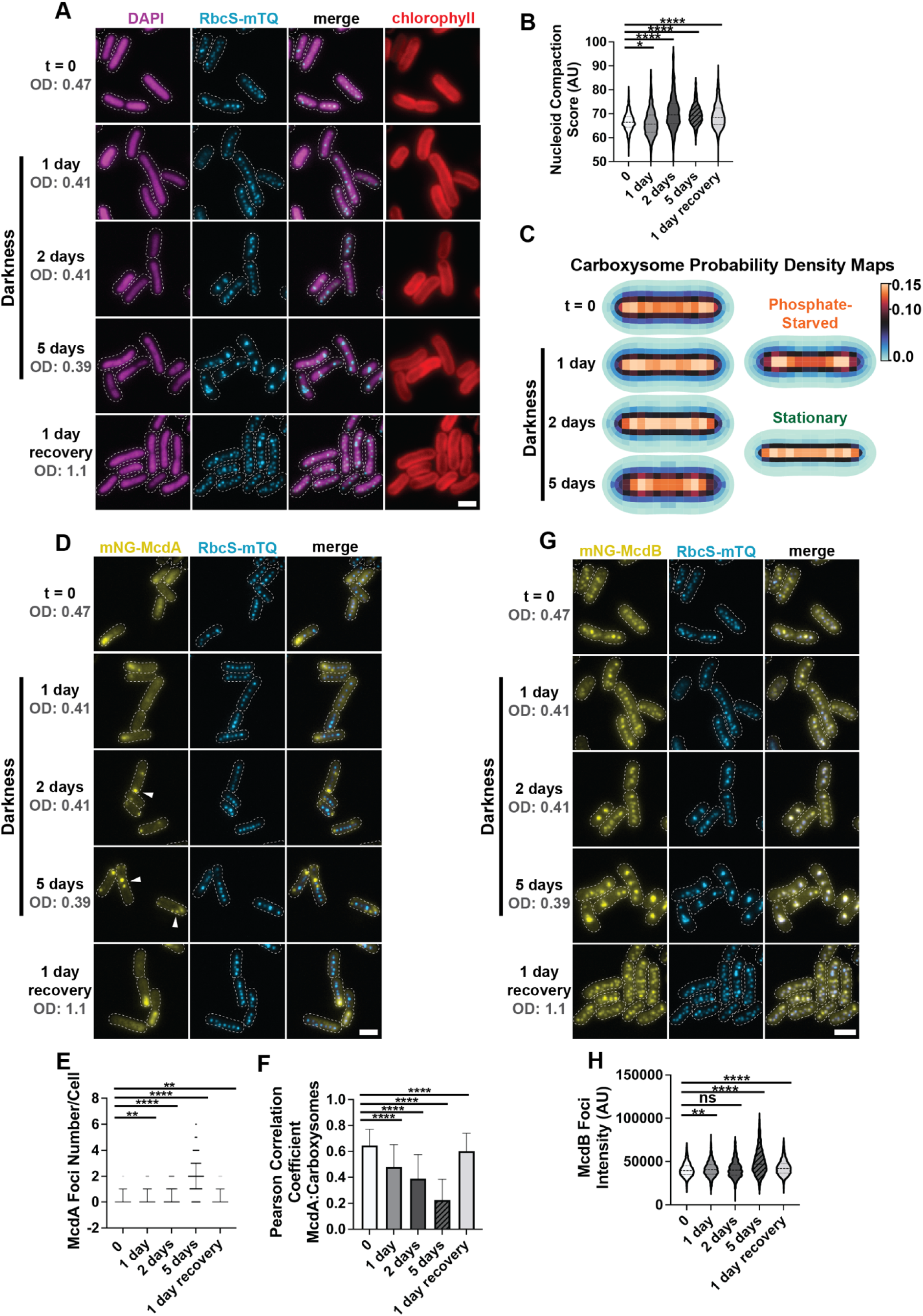
McdA is sequestered and carboxysomes cluster in prolonged darkness. **(A)** Microscopy images of DAPI-stained nucleoids (magenta) and carboxysomes (cyan) in cells grown in light deprivation conditions over 5 days and 1 day continuous light recovery. Dotted white line shows cell boundaries from phase contrast. Chlorophyll channel is red. **(B)** Nucleoid compaction score in arbitrary units. Significance from Kruskall-Wallis and Dunn’s multiple comparisons, P < 0.0001, t = 0, n = 4 biological replicates; day 1 – 5 and 1 day recovery, n = 6 biological replicates per time point, >200 cells per replicate. **(C)** Carboxysome probability density map from cells grown in prolonged darkness. t = 0, n = 840 cells; 1 day, n = 845 cells; 2 days, n = 376 cells; 5 days, n = 736 cells; 1 day recovery, n = 1961 cells; 3 days phosphate starvation, n = 1317 cells; stationary, n = 1401 cells **(D)** Microscopy images of mNG-McdA and carboxysomes (cyan) in cells grown in light deprivation conditions over 5 days and 1 day continuous light recovery. Dotted white line shows cell boundaries from phase contrast. White arrows highlight non-oscillatory foci. **(E)** mNG-McdA foci number per cell. Significance from Kruskall-Wallis and Dunn’s multiple comparisons, P < 0.0001, t = 0, n = 2 biological replicates; day 1 – 5 and 1 day recovery, n = 3 biological replicates per time point, >200 cells per replicate. **(F)** Pearson correlation coefficient from McdA and carboxysome fluorescence channels. Significance from Kruskall-Wallis and Dunn’s multiple comparisons, t = 0, n = 2 biological replicates; day 1 – 5 and 1 day recovery, n = 3 biological replicates per time point, >200 cells per replicate. **(G)** Microscopy images of mNG-McdB and carboxysomes (cyan) in cells grown in light deprivation conditions over 5 days and 1 day continuous light recovery. Dotted white line shows cell boundaries from phase contrast. **(H)** McdB foci intensity. Significance from Kruskall-Wallis and Dunn’s multiple comparisons, t = 0, n = 2 biological replicates; day 1 – 5 and 1 day recovery, n = 3 biological replicates per time point, >200 cells per replicate. (scale bar 2 μm).

The data show that prolonged-darkness halts *S. elongatus* growth, but this effect is not coupled to the drastic changes in nucleoid compaction we observed in stationary phase or phosphate-starved cells.

Prolonged-darkness therefore provided an opportunity to observe carboxysome organization in metabolically dormant cells without a highly compacted nucleoid. Unlike in stationary phase or phosphate-starved cells, carboxysome foci did not align down the medial axis of cells in prolonged-darkness (Figure 3A & C). Instead, carboxysomes clustered into fewer (Figure S3B), high-intensity foci (Figure S3C) that were more distantly spaced from one another (Figure S3D). Strikingly, carboxysomes redistributed within one day of light reintroduction (Figure 3A, Figure S3B-D). The data show that prolonged-darkness results in carboxysomes forming mispositioned clusters similar to cells lacking a functioning McdAB system.

Since the data suggested that the McdAB system is inactive, we imaged McdA and McdB localization under continuous darkness. Cellular levels of McdA declined by 25% after five days in darkness (Figure S3E), and intriguingly, McdA oscillations transitioned into punctate foci, which coincided with carboxysomes forming mispositioned clusters in the cell (Figure 3DE). The McdA foci were static (Figure S3F) and did not colocalize with mispositioned carboxysomes (Figure 3F). McdB, on the other hand, remained associated with the mispositioned carboxysome clusters over prolonged-darkness (Figure 3GH). One day of light reintroduction was sufficient to recover both McdA oscillation and carboxysome redistribution across the cell population (Figure 3 and Figure S3). Together, we conclude that under prolonged-darkness, McdA is sequestered into foci and the nucleoid remains moderately expanded, which allows carboxysomes to diffuse and cluster in the absence of active positioning by the McdAB system.

### Drug-induced nucleoid compaction overrides the McdAB system in maintaining carboxysome distribution

Nucleoid compaction is one of several physiological changes that occur when bacterial growth conditions are altered. Therefore, the data presented thus far is correlative—showing a strong correlation between nucleoid compaction and the maintenance of carboxysome distribution in the absence of active positioning by the McdAB system. We next set out to determine if drug-induced nucleoid compaction directly influenced carboxysome distribution. We have previously shown that the gyrase inhibitor, ciprofloxacin, compacts the nucleoid in *S. elongatus* ^25^. We find here that when exponential cells were treated with ciprofloxacin, carboxysomes remained distributed over hyper-compacted nucleoids in viable cells (Figure 4A, Figure S4A-C). The term ‘hyper-compacted’ is used to emphasize that the degree of nucleoid compaction is significantly greater than that in stationary phase cells (Figure 4B). Additionally, the hyper-compacted nucleoids were largely compacted along the long axis of cells, whereas nucleoids were compacted along the short axis in our nutrient deprivation experiments (Figure 4C). This hyper-compaction of the nucleoid resulted in carboxysomes being positioned further away from the medial axis of the cell (Figure 4D). Together, the data shows that nucleoid compaction influences the distribution of carboxysomes.

**Figure 4.**
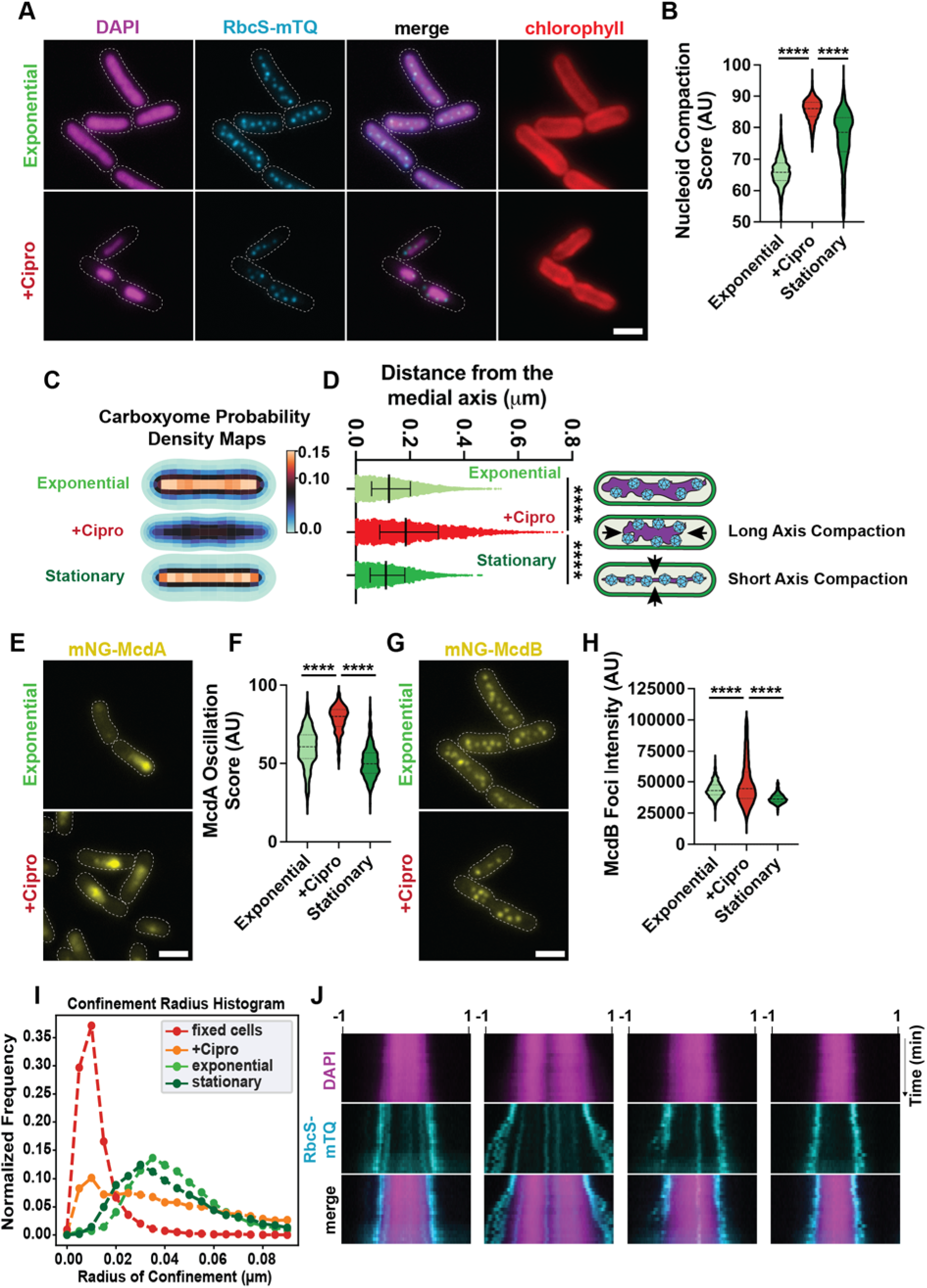
Drug-induced nucleoid compaction overrides the McdAB system in maintaining carboxysome distribution. **(A)** Microscopy images of DAPI-stained nucleoids and carboxysomes (cyan) in exponential growth with or without 50 μM Ciprofloxacin for 4 hours. Red is the chlorophyll channel. **(B)** Nucleoid compaction score in arbitrary units. Significance from Kruskall-Wallis and Dunn’s multiple comparisons, P < 0.0001, (exponential) n = 6 biological replicates, (cipro) n = 6 biological replicates, and (stationary) n = 5 biological replicates, >200 cells per replicate. **(C)** Carboxysome probability density maps from cells in exponential phase, exponential cells treated with ciprofloxacin, and stationary phase. Exponential, n = 1662 cells; ciprofloxacin, n = 969 cells; stationary, n = 1401 cells. **(D)** mTQ Foci Distance from the Medial Axis. Distances of mTQ foci from the long axis of the cell, computed using particle mapping code. Significance from Kruskall-Wallis and Dunn’s multiple comparisons, P < 0.0001, (exponential) n = 6 biological replicates, (cipro) n = 6 biological replicates, and (stationary) n = 5 biological replicates, >200 cells per replicate. **(E)** Microscopy images of mNG-McdA in cells with and without ciprofloxacin treatment. **(F)** McdA Oscillation score demonstrating oscillatory (higher score) or diffuse nature (lower score) in arbitrary units. Significance from Kruskall-Wallis and Dunn’s multiple comparisons, P < 0.0001, (exponential) n = 3 biological replicates, (cipro) n = 3 biological replicates, and (stationary) n = 2 biological replicates, >200 cells per replicate. **(G)** Microscopy images of mNG-McdB in cells with and without ciprofloxacin treatment. **(H)** McdB foci intensity. Significance from Kruskall-Wallis and Dunn’s multiple comparisons, P < 0.0001, (exponential) n = 3 biological replicates, (cipro) n = 3 biological replicates, and (stationary) n = 3 biological replicates, >200 cells per replicate. **(I)** Confinement radius histogram illustrating carboxysome foci confinement radii in exponential, exponential treated with ciprofloxacin, stationary, and fixed cells based on TrackMate analysis. **(J)** Representative kymographs of DAPI-stained nucleoids and carboxysomes (cyan) in Ciprofloxacin treated cells. Long-axis is cell length, short-axis is time, 0 to 60 minutes. (scale bar 2 μm).

We next set out to determine the localization and dynamics of the McdAB system in cells with hyper-compacted nucleoids. Interestingly, McdA displayed robust oscillation over the shortened nucleoid region of the cell (Figure 4EF, Figure S4D, **Video 5**). McdB also remained localized to carboxysomes (Figure 4G), albeit with a broader intensity distribution (Figure 4H); likely due to space limitations for distributing carboxysomes on a hyper-compacted nucleoid. Despite McdA oscillation and McdB colocalization with carboxysomes being relatively unperturbed, carboxysome dynamics were severely restricted on hyper-compacted nucleoids; more so than on the compacted nucleoids of stationary phase cells (Figure 4I). The findings show that a hyper-compacted nucleoid can override a functional McdAB system to maintain carboxysome organization in the cell. Consistent with this interpretation, carboxysomes near the poles of hyper-compacted nucleoids became more dynamic over time as polar regions of the nucleoid re-expanded following drug removal (Figure 4J).

## DISCUSSION

The nucleoid serves as a critical matrix for positioning diverse cellular cargos by ParA family ATPases, including chromosomes, plasmids, and an array of protein-based organelles, many of which are fundamental to cell survival and pathogenesis ^2,23,32,33^. Yet it remains unclear how the compaction state of the nucleoid influences the subcellular organization of these mesoscale cargos. In this study, we investigated the mechanisms underlying carboxysome organization in *S. elongatus* under physiological states that influence nucleoid compaction, particularly during stationary phase and under environmental stresses such as phosphate deprivation and prolonged-darkness. We found that, while carboxysomes remain distributed in stationary phase cells, their spatial arrangement is altered, becoming more tightly aligned along the medial axis with restricted diffusion. Surprisingly, this distribution was maintained despite the loss of McdA oscillation and the reduced colocalization of McdB with carboxysomes, suggesting that factors other than the McdAB system contribute to carboxysome positioning during stationary phase. We further demonstrated that nucleoid compaction, induced by phosphate starvation or drug treatment, correlates with carboxysome immobilization and alignment along the axis of nucleoid compaction, while prolonged-darkness led to McdA sequestration and carboxysome clustering on an expanded nucleoid (Figure 5). Together, our findings reveal that nucleoid structure plays a significant role in carboxysome positioning, particularly under conditions where the McdAB system is downregulated or inactivated.

**Figure 5.**
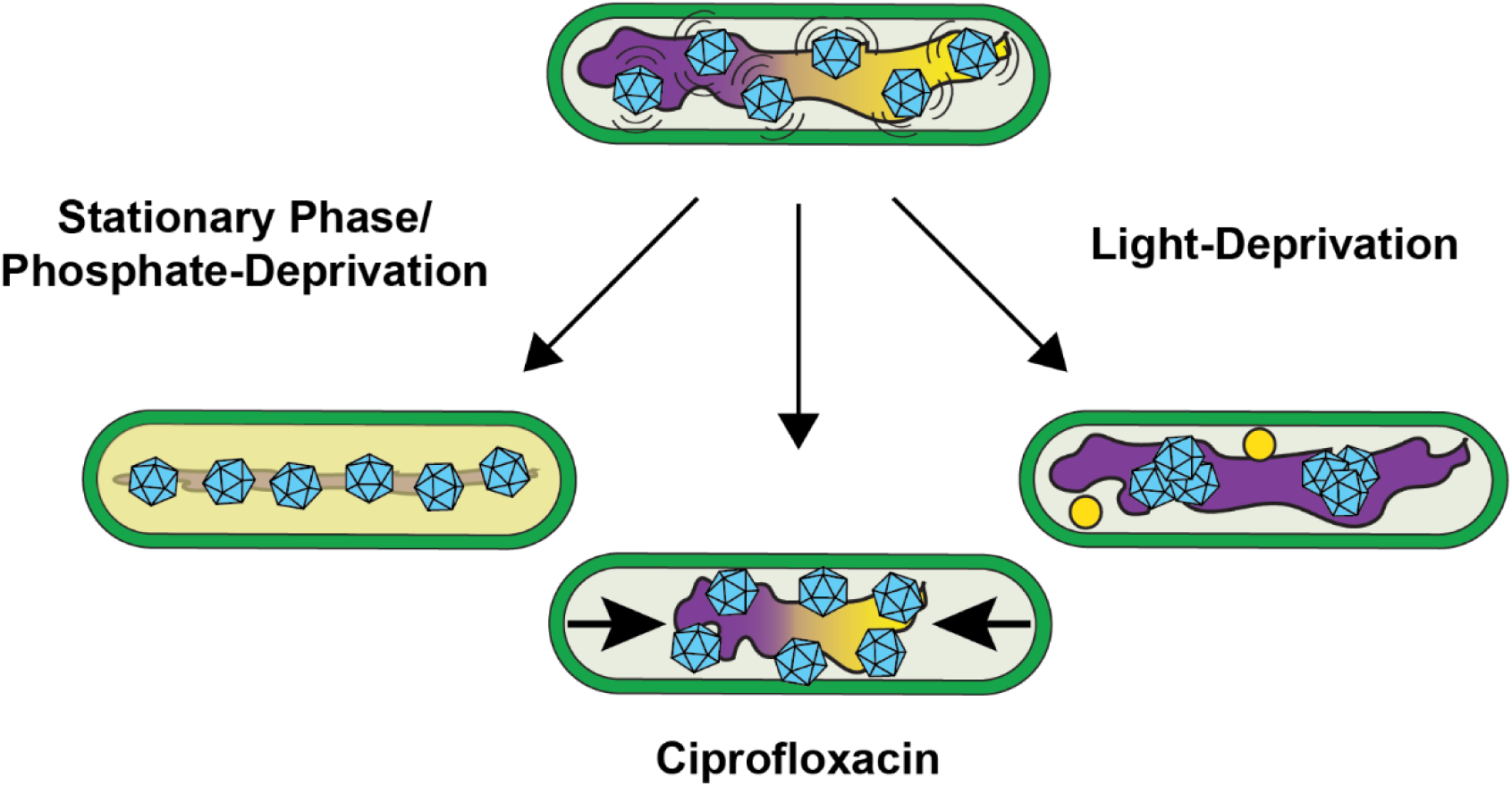
The compacted nucleoid can maintain carboxysome distribution. McdA oscillation (*yellow*) on the nucleloid (*magenta*) distributes McdB-bound carboxysomes (*cyan*) in exponential phase cells (*top*). In stationary phase or upon phosphate depletion (*left*), static carboxysomes are tightly aligned along the medial axis, despite a loss of McdA oscillation. Cirpofloxacin treatment resulted in hypercompacted nucleoids that immobilized carboxysomes, which were unresponsive to McdA oscillation (*bottom*). With light-deprivation (*right*), McdA was sequestered, resulting in carboxysomes clustering on an expanded nucleoid.

### Carboxysome distribution and suppression of McdAB in metabolically dormant cells

Our results support and extend previous observations that carboxysomes remain distributed in stationary phase cells ^30^. However, in contrast to exponential phase cells, where McdAB actively positions carboxysomes ^14,24,26^, we find that McdA loses its dynamic oscillation and McdB no longer strongly colocalizes with carboxysomes in stationary phase. Despite these differences, carboxysomes remain evenly distributed along the cell length, albeit in a more compacted linear arrangement. The data implicate the compacted nucleoid in maintaining carboxysome positioning when McdAB activity is diminished.

The decreased function and stability of McdA and McdB proteins may stem from the metabolic shifts associated with stationary phase. While McdA and McdB levels only slightly decreased, their functional impairment and mislocalization in the cell suggests regulatory control at the post-translational level. One possible explanation for McdAB inactivation in metabolically dormant cells is a stress-induced drop in cytoplasmic pH. Many bacterial stress responses, including those triggered by nutrient deprivation and energy depletion, lead to cytoplasmic acidification, which can alter protein structure, stability, and function ^37^. Given that McdA requires ATP binding for dimerization and nucleoid association, a reduction in cytoplasmic pH could directly impact its ATPase activity, oligomeric state, and nucleoid-binding ability. Similarly, McdB may undergo conformational changes that reduce its affinity for carboxysomes, further disrupting McdAB-mediated positioning.

A decrease in the cellular ATP pool is another plausible mechanism for McdA inactivation. ATP depletion is a well-documented consequence of prolonged starvation and energy stress in bacteria ^38–40^. Since McdA requires ATP binding to dimerize and interact with the nucleoid, a reduction in ATP availability could prevent the formation of active McdA dimers, thereby abolishing oscillations on the nucleoid. Our observation that McdA oscillations gradually dampen and become undetectable during phosphate starvation, or after prolonged-darkness, is consistent with a model in which ATP depletion leads to McdA inactivation.

Interestingly, McdA also formed foci in darkness that were static and did not colocalize with mispositioned carboxysomes. It is possible that, in the absence of sufficient ATP, McdA may misfold or aggregate into inactive assemblies. This type of ATP-dependent aggregation has been observed for other ATPases under energy-limited conditions and may serve as a protective mechanism to prevent protein degradation during prolonged-darkness stress ^41–43^.

The rapid recovery of McdA oscillation and carboxysome redistribution upon re-exposure to light suggests that once ATP levels are restored, McdA aggregates dissolve, allowing McdA dimers to resume function.

### Nucleoid compaction as a driver of carboxysome organization

Our findings indicate that nucleoid structure plays a key role in maintaining carboxysome organization, particularly in the absence of active McdAB positioning. We observed that in stationary phase, the nucleoid significantly compacts along the medial axis of the cell, and this compaction correlated with the linear alignment and restricted diffusion of carboxysomes (Figure 5). Similarly, phosphate starvation, which induced nucleoid compaction, resulted in carboxysomes adopting a comparable medial alignment. Together, the evidence suggests that as the nucleoid compacts, it passively restricts carboxysome movement, effectively maintaining their spatial distribution without the need for McdAB activity.

Further supporting this idea, we found that cells undergoing partial nucleoid compaction due to phosphate starvation exhibited distinct spatial patterns—carboxysomes aligned within compacted regions, while clustering was observed in expanded areas. Additionally, ciprofloxacin-induced nucleoid hyper-compaction resulted in carboxysomes being positioned further from the medial axis, highlighting the influence of nucleoid topology on carboxysome organization. These observations demonstrate that nucleoid architecture is a significant determinant of carboxysome positioning in *S. elongatus*, particularly under conditions where the McdAB system is inactive.

### The effects of prolonged darkness on carboxysome positioning

In contrast to nutrient limitation, prolonged darkness led to a distinct response in carboxysome organization. The moderate nucleoid compaction observed in darkness did not appear sufficient to maintain carboxysome distribution as seen in phosphate starvation or stationary phase. Instead of remaining evenly distributed, carboxysomes clustered into fewer, high-intensity foci—a phenotype found when the McdAB system is nonfunctional ^14,24,25,27,28^.

These findings suggest that prolonged darkness induces a metabolic state that disrupts McdAB function, leading to carboxysome mispositioning and clustering. This mispositioning was accompanied by McdA sequestration into static foci. Interestingly, upon re-exposure to light, both McdA oscillations and carboxysome distribution were rapidly restored, demonstrating the dynamic nature of these changes. The findings suggest that while a compacted nucleoid can serve as a passive organizing force, carboxysome distribution in the cell requires the McdAB system when the nucleoid is expanded.

### Nucleoid compaction can override the McdAB system

Our results with ciprofloxacin treatment provide strong evidence that nucleoid compaction can override active carboxysome positioning by McdAB. Despite McdA continuing to oscillate and McdB remaining associated with carboxysomes, carboxysome diffusion was severely restricted on hyper-compacted nucleoids. This demonstrates that nucleoid structure alone can dictate carboxysome positioning, even when McdAB remains functional.

Furthermore, we observed that as the nucleoid re-expanded following drug removal, carboxysome dynamics recovered, particularly at the expanding nucleoid poles. This suggests that nucleoid expansion allows carboxysomes to regain mobility and dynamic positioning by the McdAB system. These findings emphasize the hierarchical relationship between nucleoid structure and McdAB-mediated positioning, where nucleoid compaction can act as a dominant factor in determining carboxysome organization under specific conditions.

### Conclusions and Future Directions

Together, our findings reveal a complex interplay between active and passive mechanisms governing carboxysome organization in *S. elongatus*. While the McdAB system is essential for positioning carboxysomes in exponential growth on expanded nucleoids, our data indicate that nucleoid compaction can serve as an alternative mechanism for maintaining their distribution in the absence of a functional McdAB system. These findings raise several important questions. How is the McdAB system inactivated under different stress conditions? What molecular mechanisms link nucleoid compaction to carboxysome positioning? Future studies investigating the regulatory pathways controlling McdAB function and the biophysical properties of nucleoid-compaction-driven carboxysome organization will provide valuable insights into bacterial organelle positioning strategies.

ParA-mediated partitioning of F plasmids has been shown to highly correlate with dense regions of the bacterial nucleoid ^44^, which is consistent with the nucleoid compaction state providing an additional layer of spatial regulation. Given the shared mesoscale sizing of the diverse cellular cargos positioned by ParA family ATPases, it is highly likely that our findings extend beyond carboxysomes and apply more broadly to other macromolecular complexes and organelles that rely on the bacterial nucleoid as a matrix for spatial organization in the cell.

## MATERIALS AND METHODS

### Constructs Used

| Strain | Identifier | Description | Reference |
| --- | --- | --- | --- |
| mNG-McdA + RbcS-mTQ | C5/AH3 | mNG-McdA at native locus with native promoter, RbcS-mTQ at neutral site I with native promoter | <sup>14</sup> |
| mNG-McdB + RbcS-mTQ | C28/AH17 | mNG-McdB at native locus with native promoter, RbcS-mTQ at neutral site I with native promoter | <sup>14</sup> |

### Growth Conditions

All strains were grown in BG11 media pH 8.3 (Sigma). Cultures were grown at a 50 ml volume in 125 ml beveled flasks (Corning) or at 100 ml volume in 250 ml beveled flasks. Strains were grown in a Minitron incubator (Infors-HT) at 30L, 2% CO_2_, shaking at 130 RPM, with 60 μmol m^-2^ s^-1^ LED light. Cultures were maintained by back diluting regularly into fresh BG11 media.

### Growth Phase

Stationary phase cultures were created by allowing an exponential culture to continue to grow for several weeks without back dilution. OD_750nm_ > 10 was considered stationary phase based on our growth curve.

### Nutrient Deprivation

For phosphate deprivation conditions, K_2_HPO_4_ was removed from the BG11 media. Nutrient deprivation cultures were started from 100 ml exponentially growing cultures that were centrifuged at 4000 RPM for 10 min. After decanting the supernatant, pellets were resuspended in residual media, ∼1 m BG11. 100 - 200 μl of the resuspended pellet was then added to fresh BG11 media or BG11 media lacking phosphate to reach an initial OD_750nm_ of 0.1 - 0.2. At each time point, 2 ml of the culture was removed for imaging. Each strain was grown in triplicate and the error bars in the data represent the standard deviation across three flasks. Images are representative across all biological replicates.

### Light Deprivation

Light deprivation cultures were started from 50 ml exponentially growing cultures grown in continuous light. The cultures were split into two flasks and diluted with 25 ml fresh BG11 to reach an initial OD_750nm_ of 0.4 - 0.5. Cultures grown in darkness had no light source and the incubator window was blocked with cardboard. At each time point, 2 ml of the culture was removed for imaging. Cultures were recovered by transferring them back to continuous light conditions for one day. Experiments were performed twice and across three biological replicates. Images are representative of all biological replicates.

### Ciprofloxacin Treatment

25 ml of cells in a 125 ml beveled flask were treated with 50 μM ciprofloxacin for 4 hours to induce nucleoid compaction. To visualize the compacted nucleoid region, 2 ml of ciprofloxacin-treated *S. elongatus* cells were harvested by centrifugation at 16,000 × g for 1 min. The pelleted cells were then washed and resuspended in 100 μl of phosphate-buffered saline (pH 7.2).

### Growth Curve Measurements

Growth curve cultures were started from 100 ml exponentially growing cultures that were centrifuged at 4000 RPM for 10 min. After decanting the supernatant, pellets were resuspended in residual media, ∼1 ml BG11. 100 - 200 μl of the resuspended pellet was then added to fresh BG11 media to reach an initial OD750_nm_ of 0.1 - 0.2. At each time point, the OD750_nm_ was measured using a DS-11 spectrophotometer (Denovix). Each strain was grown in triplicate with the error bars representing the standard deviation across the 3 flasks.

### DAPI Staining

2 ml of culture was centrifuged at 16,000 × g for 1 min. Cells were then washed with 1X PBS (pH 7.2) and then resuspended in 100 ul of PBS. DAPI (8 μl from a 20 μg/ml stock concentration) was added to the cell suspension followed by 20-min incubation in the dark at 30°C, 2% CO_2_, and shaking at 130 RPM. DAPI-stained cells were washed twice with 1 ml H2O and then resuspended in 100 μl H2O prior to visualization using the DAPI channel.

### Live Fluorescence Microscopy

After DAPI staining, 1 μl of cells were plated on a disc made of 1.5% UltraPure agarose (Invitrogen) in BG11, which was then placed on a 35-mm cover glass-bottom dish (MatTek). Imaging was performed through the NIS Elements software on a Nikon Ti2-E inverted microscope with a 100X objective lens, SOLA LED light source, and a Hamamatsu camera. mNG-McdA and mNG-McdB were imaged using a YFP filter, RbcS-mTQ carboxysomes with a CFP filter, and DAPI stained DNA was visualized with a DAPI filter.

### Data Analysis

Outliers were removed from all graphs using the ROUT method in GraphPad Prism.

### Image analysis

#### Confinement radius

The first ten localizations of each trajectory were used to calculate the confinement radius. The confinement radius was determined by finding the average xy position of the cropped track by averaging the track coordinates. The average of the difference between this position and the track coordinates gives the confinement radius.

#### Cell registration

Phase contrast images were registered to the first frame in the stack using cross-correlation. If the stack contained multi-channel data, the shifts used for the phase contrast images were applied to all other time-corresponding frames in other channels. The registered stacks were then saved as separate TIFF stacks.

#### Cell segmentation

Drift-corrected phase contrast images were used for cell segmentation via the Omnipose ^45^ package in Python. A custom Python script was implemented for batch segmentation: *phasemasks_omni.py*. Prior to segmentation, a Gaussian blur (standard deviation of Gaussian = 66 nm) was applied to the images. The ‘bact_phase’ pre-trained model was used for segmentation. Cells touching the image borders were ignored, and erroneous segmentations were manually corrected using the Omnipose GUI or excluded from further analysis if cell boundaries were not clear from the phase contrast image. Some cell masks had gaps within the inner region of the cell representing the chlorophyll localization pattern. We corrected these errors using the binary_fill_holes function from the scipy.ndimage package ^46^. Resulting masks were used for subsequent single-cell analyses. For cell morphology analysis, the cell mask was used to determine the cell length, width, and area.

#### Fluorescent focus detection

Carboxysome, McdA, and McdB foci were detected using a custom Python script. First, the cell masks were used to individually analyze every cell in the field of view. The fluorescence signal in each cell was then sharpened using the scikit-image ^47^ function ‘filters.unsharp_mask’; this step was followed by a Gaussian blur. Using this pre-processed image, fluorescent foci were detected with the Laplacian of Gaussian (LoG) algorithm. Parameters used for these preprocessing steps can be found in Supplementary Table 1.

To calculate the intensity of a fluorescent focus, the focus area was determined by the resulting coordinates and radius of the LoG detection. The area of the focus was defined by a circle centered on the xy coordinate of the detection with radius = 2*r*, with *r* being the LoG determined spot radius. The pixel intensities within this circle were summed to define the focus intensity.

#### Carboxysome localization heatmaps

The localization coordinates of detected carboxysome foci and cell masks were used to make localization density heatmaps with the *spideymaps* tool (https://github.com/BiteenMatlab/spideymaps) in Python. For each segmented cell, morphological skeletonization was used to define the midline bisecting the cell lengthwise.

Inspired by the *colicoords* package ^48^, the cartesian coordinates of each foci (x, y) were re-parameterized into a three coordinate system: *r, l*, and *ϕ*. *r* is the radial distance from the midline. l is the longitudinal distance along the length of the cell found from projection onto the midline. *ϕ* is the angle meaured relative to the midline in the polar regions. r and l were divided by the average cell width and total length, respectively, to calculate *r_rel_* and *l_rel_* so that all localizations share a common frame of reference. Localizations were binned based on *r_rel_, l_rel_*, and *ϕ*. Because this approach produces unequal bin areas, bin areas were calculated for each bin in every cell. Localization counts and associated bin areas were summed across all cells.

Areas and counts were symmetrized fourfold by adding counts and areas from symmetrically equivalent bins. The summed localization counts were divided by the summed areas to calculate average localization densities (counts per pixel^2^). Count densities were converted to probability densities by dividing by the total number of counts across all cells and multiplying by the number of cells. Probability density maps are reported with units of µm^-2^.

To generate representative cells for visualization, masks for individual cells were aligned and summed. 50% of the max of summed masks was used as a threshold to create representative masks. The midlines and outlines of the representative masks were used to create the bin shapes for the figures.

#### McdA distribution

To analyze the localization of mNG-McdA in single cells, we implemented the same pixel intensity distribution analysis used previously ^49^. Briefly, the intensity of all the pixels in a cell were normalized by the minimum and maximum pixel intensity values in the corresponding cell:

*I_n_* = (*I*−*I_min_*)/(*I_max_*−*I_min_*). These normalized pixel intensities were then binned to generate histograms that represent the localization pattern of the protein. To calculate the oscillation score we determined the fraction of pixels with a normalized intensity below a threshold value, *I_n_*<L0.5 for each cell.

Drift-corrected time-lapse image stacks were used to generate kymographs along the cell length. In Fiji ^50^, a line was manually drawn along the long axis of a cell. The multi-kymograph tool was used to extract the intensity profile with a 11-pixel (660 nm) width. For visualization, the intensity profile at each time point of the kymograph was normalized to minimum and maximum intensity values within the corresponding time point. This post-processing removed the effects of photobleaching in the visualization of mNG-McdA.

#### Nucleoid morphology

To determine the nucleoid region within the cell, the cell mask was used to determine the pixel intensities within the inner rim of the cell boundary. The distribution of the inner rim pixel intensities was fit to a Gaussian distribution to determine the average intensity of the cytoplasmic region of the cell. This intensity was subtracted across all pixels in the cell. A Gaussian blur (standard deviation of Gaussian = 66 nm) was applied to the background subtracted image. Next, the nucleoid region was detected using Yen’s thresholding ^51^. The pixels in the resulting region were considered as the nucleoid. To calculate the nucleoid compaction score, *s_comp_*, we subtracted the ratio of nucleoid area to total cell area from 1: *s_comp_* = 1 – (*A_nuc_* / *A_cell_*).

## ACKNOWLEDGEMENTS

We would like to thank the members of the Vecchiarelli Lab for providing feedback on experimental design and data visualization. This work was supported by funding from the National Science Foundation to A.G.V. (Grant no. 1941966) and the National Institute of General Medical Sciences of the National Institutes of Health to A.G.V. (Grant no. R35GM152128). C.E.D. acknowledges funding from the Rackham Precandidate Research Grant.

## AUTHOR CONTRIBUTIONS

Conceptualization, C.E.D. and A.G.V.,; Investigation, C.E.D. and O.L.; Software, C.E.D, C.A.A., and D.J.F.,; supervision, A.G.V. and J.S.B.; writing – original draft, C.E.D., and A.G.V.; writing – review & editing, C.E.D., C.A.A., D.J.F., O.L., J.S.B., and A.G.V.

## Declaration of Interests

The authors declare no competing interests.

## Video Legends

**Video 1** - **Carboxysomes (blue, RbcS-mTQ) are less dynamic in stationary phase cells.** 1 hour video, each frame is 3 minutes. (scale bar 2 μm).

**Video 2** - **Carboxysomes (blue, RbcS-mTQ) in exponential phase cells**. 1 hour video, each frame is 3 minutes. (scale bar 2 μm).

**Video 3** - **McdA (yellow, mNG-McdA) robustly oscillates in exponential cells.** 1 hour video, each frame is 3 minutes. (scale bar 2 μm).

**Video 4** - **The McdA (yellow, mNG-McdA) oscillation becomes diffuse in stationary phase cells.** 1 hour video, each frame is 3 minutes. (scale bar 2 μm).

**Video 5** - **McdA (yellow, mNG-McdA) oscillates over the ciprofloxacin compacted nucleoid (magenta, DAPI)**. 1 hour video, each frame is 3 minutes. (scale bar 2 μm).

## Notes

### Competing Interest Statement

The authors have declared no competing interest.

