## Supplementary Information for "Nucleoid Compaction Influences Carboxysome Localization and Dynamics in *Synechococcus elongatus* PCC 7942"

**SUPPLEMENTAL FIGURES**

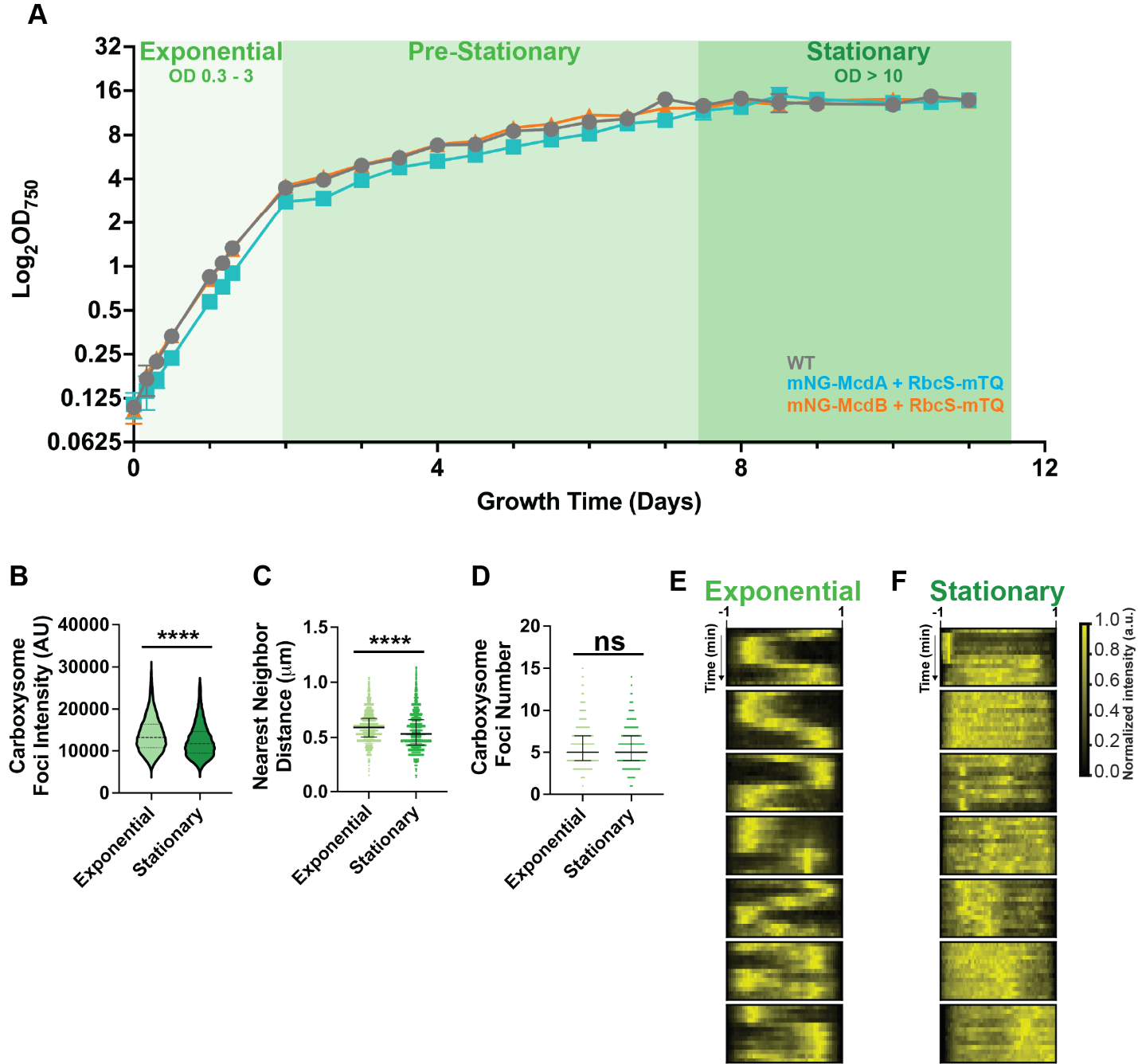

**Figure S1 - (A)** Growth curve of WT *S. elongatus* and the indicated fluorescently labeled strains. Fluorescent labeling does not significantly affect growth rate. n = 3 biological replicates per strain. Exponential phase is defined as OD_750nm_ 0.3 - 3. Stationary is defined as OD_750nm_ > 10. **(B)** Carboxysome foci intensity. Significance from Mann-Whitney test, P < 0.0001, (exponential) n = 6 biological replicates and (stationary) n = 5 biological replicates, >200 cells per replicate. **(C)** Nearest neighbor distance between carboxysomes. Significance from Mann-Whitney test, P < 0.0001, (exponential) n = 6 biological replicates and (stationary) n = 5 biological replicates, >200 cells per replicate. **(D)** Carboxysome number per cell. Significance from Mann-Whitney test, P > 0.05, (exponential) n = 6 biological replicates and (stationary) n = 5 biological replicates, >200 cells cells per replicate. **(E)** Representative kymographs of McdA dynamics in exponential phase cells. **(F)** Representative kymographs of mNG-McdA dynamics in stationary phase cells. Long-axis is cell length, short-axis is time, 0 to 60 minutes.

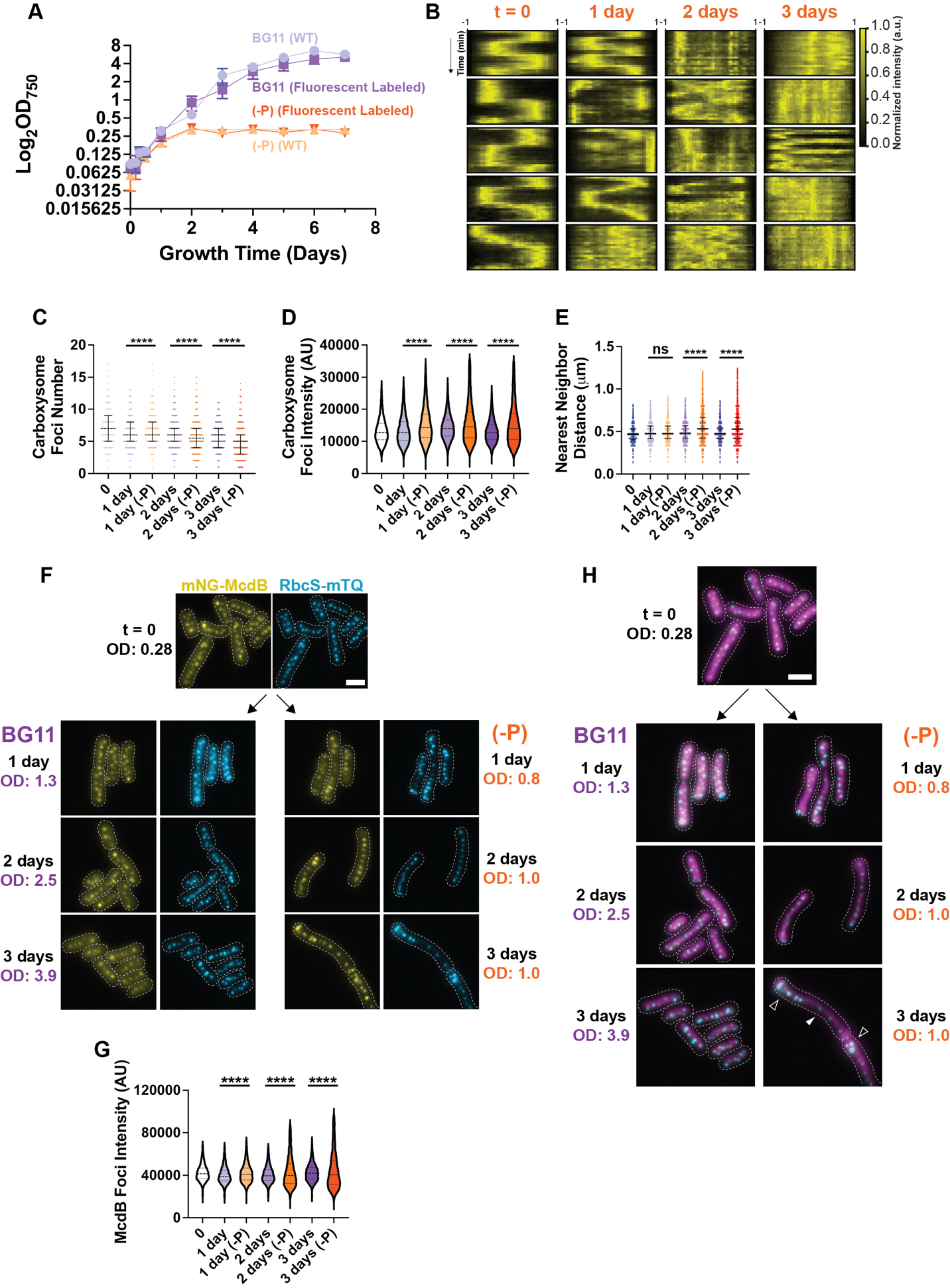
 **Figure S2 - (A)** Growth curve of WT *S. elongatus* and fluorescently labeled strains grown in BG11 and BG11 lacking phosphate (-P). Fluorescent labeling does not significantly affect growth rate. n = 3 biological replicates per treatment. **(B)** Representative kymographs of mNG-McdA dynamics in phosphate deprivation conditions over 3 days. Long-axis is cell length, short-axis is time, 0 to 60 minutes. **(C)** Carboxysome number per cell. Significance from Kruskall-Wallis and Dunn’s multiple comparisons, P < 0.0001, (BG11) n = 4 biological replicates and (-P) n = 6 replicates, >200 cells per replicate. **(D)** Carboxysome foci intensity. Significance from Kruskall-Wallis and Dunn’s multiple comparisons, P < 0.0001, (BG11) n = 4 replicates and (-P) n = 6 replicates, >200 cells per replicate. **(E)** Distance between carboxysome and its nearest carboxysome neighbor. Significance from Kruskall-Wallis and Dunn’s multiple comparisons, P < 0.0001, (BG11) n = 4 replicates and (-P) n = 6 replicates, >200 cells per replicate. **(F)** Microscopy images of mNG-McdB in cells grown in BG11 and phosphate-limited BG11 over a 3-day time course. Carboxysomes are in cyan. Dotted white line shows cells’ boundaries from phase contrast. **(G)** McdB foci intensity. Significance from Kruskall-Wallis and Dunn’s multiple comparisons, P < 0.0001, (BG11) n = 2 biological replicates and (-P) n = 3 biological replicates, >200 cells per replicate. **(H)** Merged microscopy images of DAPI stained nucleoid and carboxysomes (cyan) in cells grown in BG11 and phosphate-limited BG11 over a 3-day time course. White arrows indicate areas where carboxysomes are singly spaced on a compacted nucleoid and clumped in expanded regions. (scale bar 2 μm).

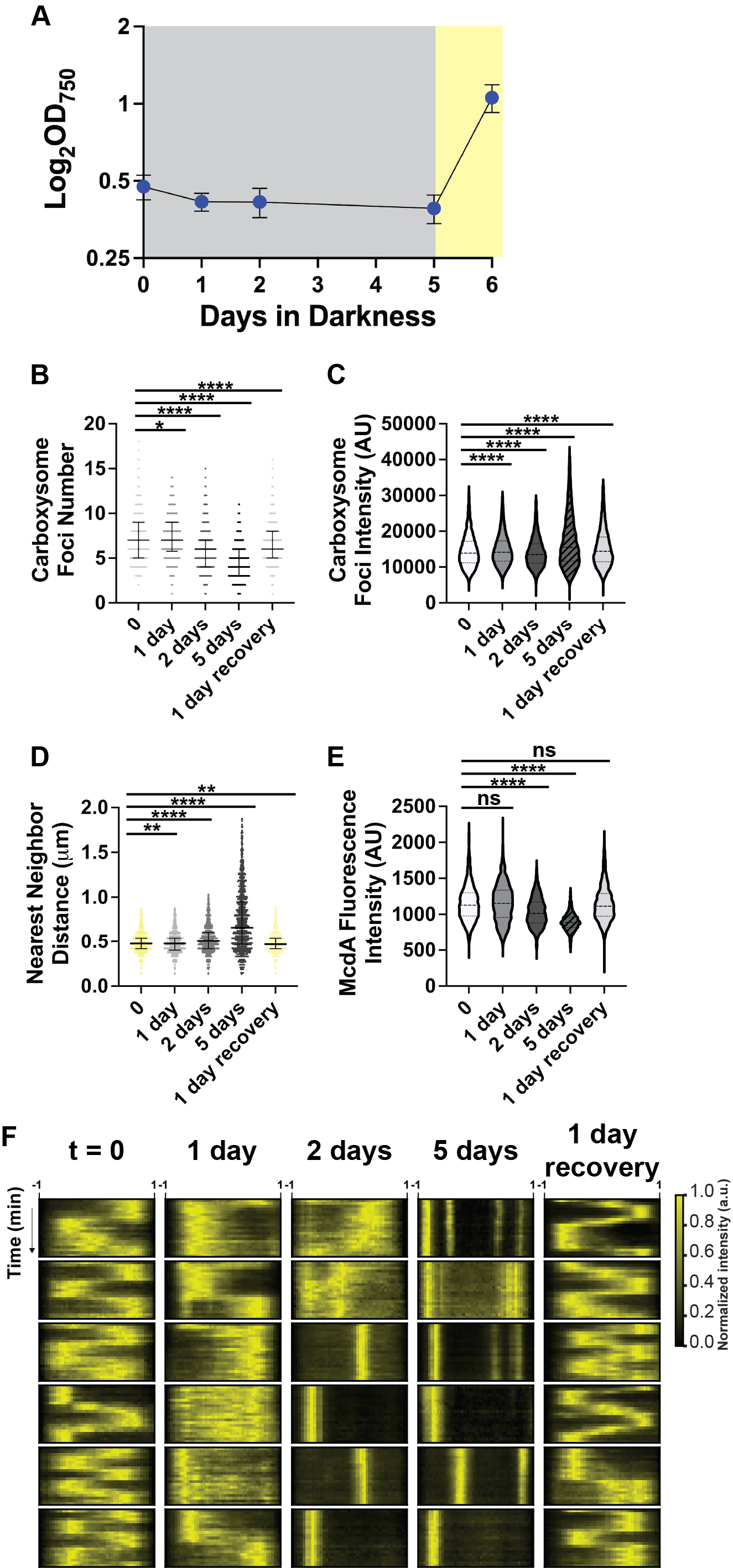

**Figure S3 - (A)** Growth curve of *S. elongatus* incubated in continuous darkness for 5 days then recovered in continuous light for 1 day. n = 4 biological replicates. **(B)** Carboxysome number per cell. Significance from Kruskall-Wallis and Dunn’s multiple comparisons, P < 0.000,1 t = 0, n = 4 biological replicates; day 1 – 5 and 1 day recovery, n = 6 biological replicates per time point, >200 cells per replicate. **(C)** Carboxysome foci intensity. Significance from Kruskall-Wallis and Dunn’s multiple comparisons, P < 0.000,1 t = 0, n = 4 biological replicates; day 1 – 5 and 1 day recovery, n = 6 biological replicates per time point, >200 cells per replicate. **(D)** Nearest neighbor distance between carboxysomes. Significance from Kruskall-Wallis and Dunn’s multiple comparisons, P < 0.000,1 t = 0, n = 4 biological replicates; day 1 – 5 and 1 day recovery, n = 6 biological replicates per time point, >200 cells per replicate. **(E)** mNG-McdA whole cell fluorescence intensity normalized by cell length. Significance from Kruskall-Wallis and Dunn’s multiple comparisons, P < 0.000,1 t = 0, n = 2 biological replicates; day 1 – 5 and 1 day recovery, n = 3 biological replicates per time point, >200 cells per replicate. **(F)** Representative kymographs of McdA dynamics in prolonged darkness conditions over 5 days and 1 day of growth recovery in light. Long-axis is cell length, short-axis is time, 0 to 60 minutes. (scale bar 2 μm).

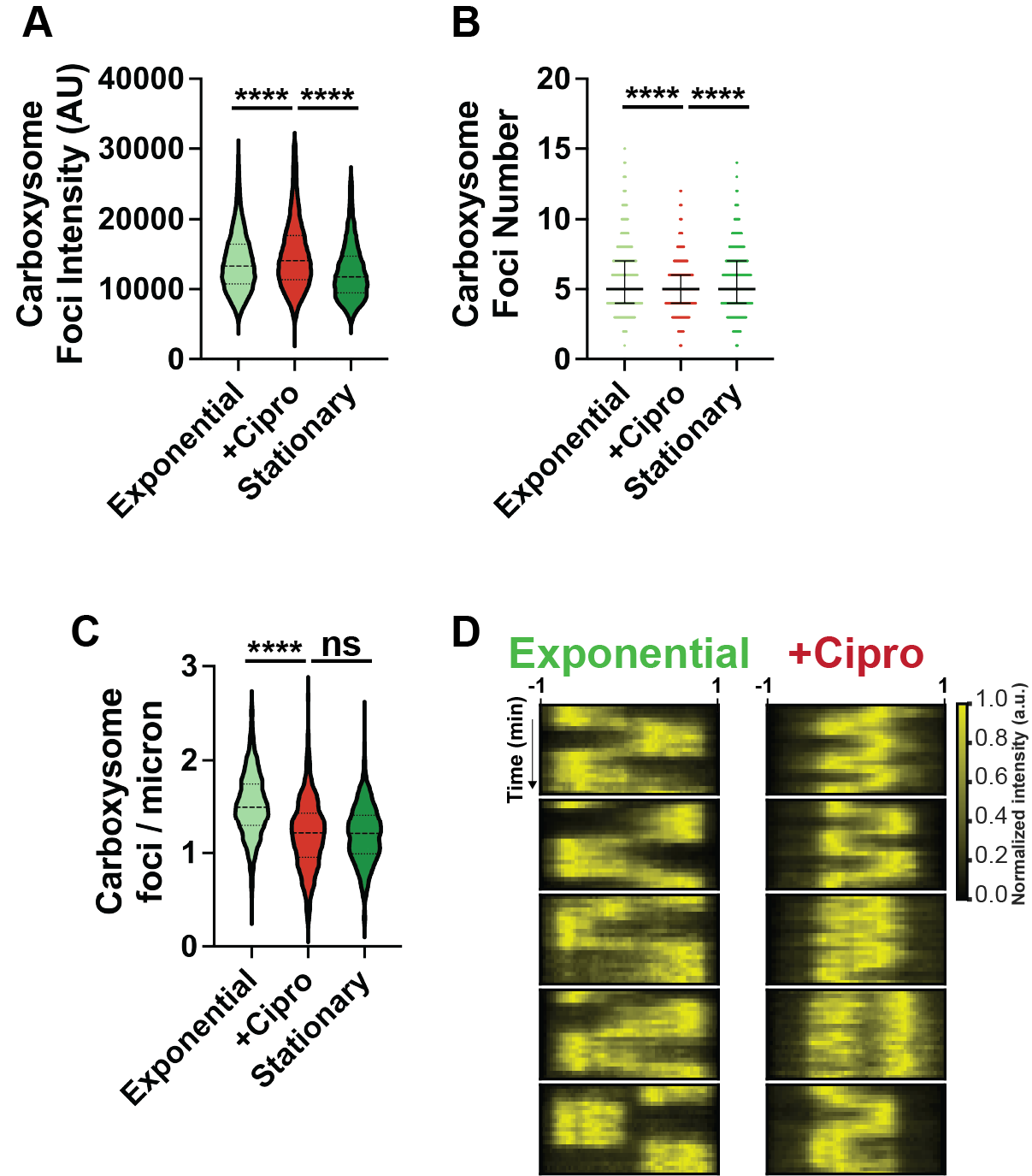

**Figure S4 - (A)** Carboxysome foci intensity. Significance from Kruskall-Wallis and Dunn’s multiple comparisons, P < 0.0001, (exponential) n = 6 biological replicates, (cipro) n = 6 biological replicates, and (stationary) n = 5 biological replicates, >200 cells per replicate. **(B)** Carboxysome number per cell. Significance from Kruskall-Wallis and Dunn’s multiple comparisons, P < 0.0001, (exponential) n = 6 biological replicates, (cipro) n = 6 biological replicates, and (stationary) n = 5 biological replicates, >200 cells per replicate. **(C)** Carboxysomes per unit of cell length, calculated by dividing the number of foci by cell length. Significance from Kruskall-Wallis and Dunn’s multiple comparisons, P < 0.0001. Significance from Kruskall-Wallis and Dunn’s multiple comparisons, P < 0.0001, (exponential) n = 6 biological replicates, (cipro) n = 6 biological replicates, and (stationary) n = 5 biological replicates, >200 cells per replicate. **(D)** Representative kymographs of McdA dyanmics in exponential phase cells with and without ciprofloxacin treatment. Long-axis is cell length, short-axis is time, 0 to 60 minutes.

**SUPPLEMENTARY TABLE 1**

**Fluorescent Focus Detection Parameters**

Carboxysomes:

| Background Subtraction | restoration.rolling_ball(cell_fluor,radius = 3) |
| --- | --- |
| Image Denoise | filters.unsharp_mask(cell_fluor_bg, radius = 2, amount = 5) |
| Add Gaussian Blur | filter.gaussian(cell_fluor_sharp, 0.01) |
| Carboxysome Detection | min_sigma = 2.75, max_sigma = 3, threshold = 0.005, overlap = 0.75 |

McdA Foci:

| Background Subtraction | restoration.rolling_ball(cell_fluor,radius = 5) |
| --- | --- |
| Image Denoise | filters.unsharp_mask(cell_fluor_bg, radius = 3, amount = 2) |
| Add Gaussian Blur | filter.gaussian(cell_fluor_sharp, 0.5) |
| McdA Foci Detection | min_sigma = 1.5, max_sigma = 5, threshold = 0.01, overlap = 0.95 |

McdB Foci:

| Background Subtraction | restoration.rolling_ball(cell_fluor,radius = 2.5) |
| --- | --- |
| Image Denoise | filters.unsharp_mask(cell_fluor_bg, radius = 1.5, amount = 1) |
| Add Gaussian Blur | filter.gaussian(cell_fluor_sharp, 0.01) |
| McdB Foci Detection | min_sigma = 1.5, max_sigma = 5, threshold = 0.003, overlap = 0.75 |
